# MetFinder: a neural network-based tool for automated quantitation of metastatic burden in histological sections from animal models

**DOI:** 10.1101/2023.09.07.555360

**Authors:** Alcida Karz, Nicolas Coudray, Erol Bayraktar, Kristyn Galbraith, George Jour, Arman Alberto Sorin Shadaloey, Nicole Eskow, Andrey Rubanov, Maya Navarro, Rana Moubarak, Gillian Baptiste, Grace Levinson, Valeria Mezzano, Mark Alu, Cynthia Loomis, Daniel Lima, Adam Rubens, Lucia Jilaveanu, Aristotelis Tsirigos, Eva Hernando

**Affiliations:** Department of Pathology, NYU Grossman School of Medicine, New York, NY 10016; Interdisciplinary Melanoma Cooperative Group, Perlmutter Cancer Center, NYU Langone Health, New York, NY 10016; Applied Bioinformatics Laboratories, NYU Langone Health, New York, NY 10016; Department of Cell Biology, NYU School of Medicine; New York, NY, USA; Experimental Pathology Research Laboratory, Division of Advanced Research Technologies, NYU Grossman School of Medicine, New York, NY 10016; Research Software Engineering Core, Medical Center Information Technology Department, NYU Langone Health, New York, NY 10016; Department of Medicine, Yale University, New Haven, CT 06510

## Abstract

Diagnosis of most diseases relies on expert histopathological evaluation of tissue sections by an experienced pathologist. By using standardized staining techniques and an expanding repertoire of markers, a trained eye is able to recognize disease-specific patterns with high accuracy and determine a diagnosis. As efforts to study mechanisms of metastasis and novel therapeutic approaches multiply, researchers need accurate, high-throughput methods to evaluate effects on tumor burden resulting from specific interventions. However, current methods of quantifying tumor burden are low in either resolution or throughput. Artificial neural networks, which can perform in-depth image analyses of tissue sections, provide an opportunity for automated recognition of consistent histopathological patterns. In order to increase the outflow of data collection from preclinical studies, we trained a deep neural network for quantitative analysis of melanoma tumor content on histopathological sections of murine models. This AI-based algorithm, made freely available to academic labs through a web-interface called MetFinder, promises to become an asset for researchers and pathologists interested in accurate, quantitative assessment of metastasis burden.

## Introduction

Mouse models represent a pivotal bottleneck in cancer biology experimentation ^1^. *In vivo* xenograft, allograft, and genetically engineered mouse models enable the testing of a hypothesis in the context of an entire mammalian organism. Rigorous *in vivo* experiment design, however, is rife with potential sources of variation. To name a few in allograft and xenograft experimental design: which cell line will be used? Which site of injection? What is the genetic background of the mice being injected? And even after all of these critical decisions are made, output data are often difficult to gather, quantify and interpret. The most prevalent method is bioluminescence imaging (BLI), in which cancer cell lines are transduced with a reporter luciferase construct ^2^. BLI is convenient for measuring tumor progression over time in living mice or *ex vivo* at sacrifice ^3–7^ but lacks resolution. When cancer cells are made to express a fluorescent reporter, *ex vivo* immunofluorescence microscopy of whole organs can give a quantitative output of tumor burden ^8–12^ but small or dim tumors may be obscured by overlying tissue. Additionally, cancer cells may be identified and counted using antibodies targeting marker genes via immunofluorescent or immunohistochemical staining of tissue sections ^4,7,13,14^. However, reporters and markers can be expressed heterogeneously and may be immune-edited out in immune competent mice ^15–17^. *In vivo* or *ex vivo* magnetic resonance imaging (MRI) is exhaustive and requires no expression of reporters, but can be prohibitively time-consuming and expensive ^3,7,18^. Histopathology allows visualization of tumor burden in cell-by-cell detail, but this time-consuming technique relies on visual recognition of reproducible patterns by trained pathologists under the microscope. Variations between observers in reporting tumor content is well known and has been reported in various studies ^19–21^.

In these ways, efforts by the cancer research community to characterize mechanisms of metastasis through xenografts, allografts, and genetically engineered animal models are hindered by limited access to pathology expertise, inter- and intra-observer variability, and lack of quantitative and high-throughput methods ^22^. Proof-of-principle studies on human tissues have revealed AI-based methods as powerful tools that leverage histological information to classify tumors based on genetic mutations ^23–26^, recurrence, death ^27,28^, or response to therapy^29^ with high accuracy. Annotations of slides have been made easier with tools like QuPath ^30^, TissueWand ^31^ or ImageJ/Fiji ^32–34^. Automatic segmentation of mouse tissues using deep learning has recently emerged, with studies of kidney diseases ^35^, lung adenocarcinoma ^36^ or lung metastatic tumors, lung fibrosis with inflammation, and fatty vacuole quantification in liver ^37–40^. However, tool development skills are needed to develop custom packages, and powerful computers with high storage and processing capabilities are required to execute such deep learning tasks. Nonetheless, such machine learning breakthroughs have not yet been applied to translational research in an approachable way, where they have the potential to expedite critical *in vivo* experiments by automating tedious quantification tasks. HALO AI (https://indicalab.com/halo-ai/) is a commercial software option that makes deep learning on histopathologic data more user-friendly, but it is not freely available for research community use.

Unsupervised image analysis and labeling can reduce dependency on pathology expertise, increase the throughput of evaluation of large whole-slide libraries and reduce the variability among observers. We therefore developed MetFinder, an online web-interface tool where researchers can upload images of their mouse tissue slides, stained only with hematoxylin and eosin, and retrieve quantitative information regarding the tumor content. The tool relies on inception v3 deep-learning architecture ^41^ re-trained for classification of tumor tiles.

Using whole-slide images annotated by board-certified anatomic pathologists, we trained the inception v3 algorithm to recognize liver and brain tissue samples from melanoma metastasis-bearing mice, resulting from xenograft or allograft transplantation. During testing, our metastasis annotating algorithm accurately marked the normal and tumor areas within any given tissue section, distinguishing them from artifacts of staining or foreign tissues across multiple xenograft models. Finally, AI-based results on an independent set of tissue sections were cross-referenced with annotations by certified pathologists, which revealed the high concordance and sensitivity of our algorithm, further supporting its value for identification and quantitative measurement of metastases in tissue sections. MetFinder even proved just as effective as manual annotation at delineating tumors from a breast cancer allograft model in the brain while requiring a fraction of the hands-on time. As a publicly available tool, MetFinder promises to standardize and accelerate the characterization of preclinical models for various cancer types.

## RESULTS

For proof-of-principle studies, human xenograft models were generated injecting melanoma cells (12273BM, 10230BM, 131/4-5B1, 113/6-4L, MeWo, SKMEL2, see **Tables S1 - S3, Figure 1**) into the left ventricles of 141 immunodeficient mice (Nod/Scid/Il2γRnull or Foxn1nu, see Table 1) using ultrasound guided intracardiac injection ^42^ during various experiments. This route of injection allows general hematogenous spread, with metastatic growths in the liver, kidney, adrenal glands, and brain depending upon the cell line used and genetic background of the recipient mice. Mice were euthanized at previously determined humane time points and tissues collected for formalin fixation and paraffin embedding following standardized protocols (Methods). Sections were collected at multiple levels and stained with hematoxylin and eosin (H&E) (Methods). Slides were scanned at 40x magnification using a Leica AT2 whole slide scanner.

**Figure 1.**
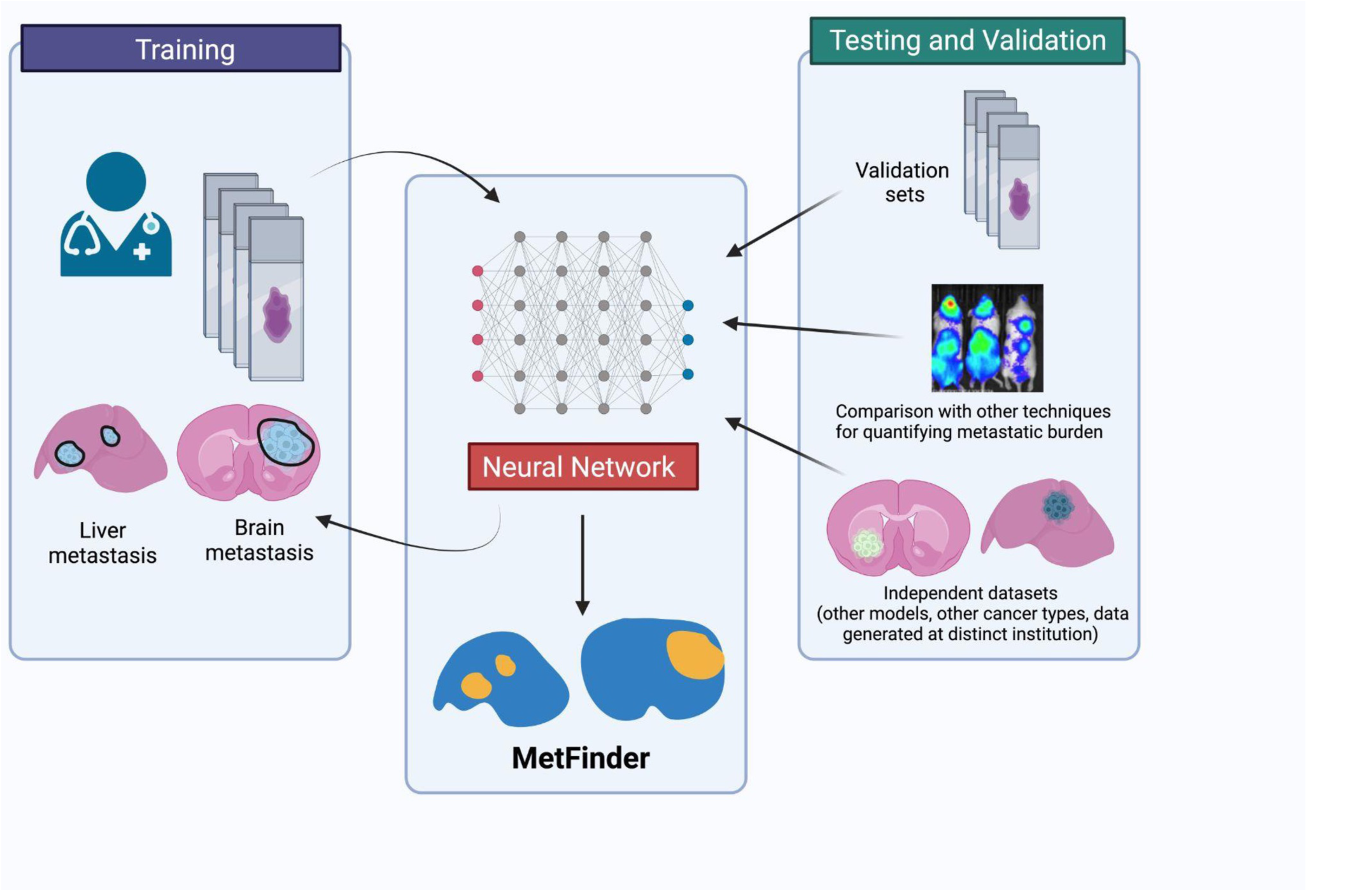
A schematic depicting the overall approach in designing MetFinder. Graphics generated using Biorender.

### Training and validation of an AI-based brain metastasis identification tool

Five independent experiments were used for initial training and validation of the brain metastasis identification tool. Training cohorts included 312 scanned H&E-stained brain slides from 141 mice harboring melanoma metastases and 14 slides from 8 negative control (no melanoma) mice. Slides were annotated by three board-certified neuropathologists (using Aperio ImageScope from Leica Biosystems or OMERO Plus from Glencoe Software), who demarcated tumor areas (Supp Figure 1). On 5-fold cross-validation, our AI-based tool displayed an AUC (Area Under the Receiver Operating Curve) of 0.997 (std=16e-4, baseline of random classifier being 0.5 for AUCs) & 0.997 (std=10e-4) and PR (Precision-Recall) of 0.984 (std=77e-4, baseline of 0.067) & 0.999 (std=3e-4, baseline of 0.035) in tumor and non-tumor identification respectively (**Figure 2A**). Using the whole training set and applied on the external dataset (**Figure 2B**), an ROC close to 1.00 is reached for tumor detection (confidence intervals CIs=0.9998-1.000), 0.997 for non-tumor regions (CIs=0.994-0.999) and 0.986 for artifacts (CIs=0.977-0.993), and a PR close to 1.00 for tumor and non-tumor (CIs=0.997-1.000 and CIs=0.999-1.000 respectively), and 0.853 for artifacts (CIs=0.784-0.916), assuming each test tile is assigned the label of the mask drawn by the expert if it overlaps with it by at least 50%. As expected, if that threshold is set to 100% to estimate the performance on regions more homogeneous (less likely to contain features from 2 different classes) and ignore the potentially heterogenous edge of the selected regions, the performance increases further (**Table S4**). The confusion matrix shows an overall specificity of 0.954, accuracy of 0.970, precision of 0.746, recall/sensitivity of 0.907 and F1-score (harmonic mean of precision and recall) of 0.761 (**Table S5)**. This performance is especially remarkable considering that the cell line giving rise to the tumors in this external dataset was not represented in the training cohort. However, this experiment was still carried out at the same institution as the training cohort, meaning the embedding, sectioning, and staining were all carried out by the same facility. With the aim of making the tool useful to labs around the world, we tested its performance on mouse brain sections generated at another institution. In addition to exhibiting tumors from cell lines the tool had not been trained on, these sections were cut on a transverse plane, distinct from the coronal view used in the training set. Further, they were stained by different hands at an institution with distinct standard operating procedures.

**Figure 2.**
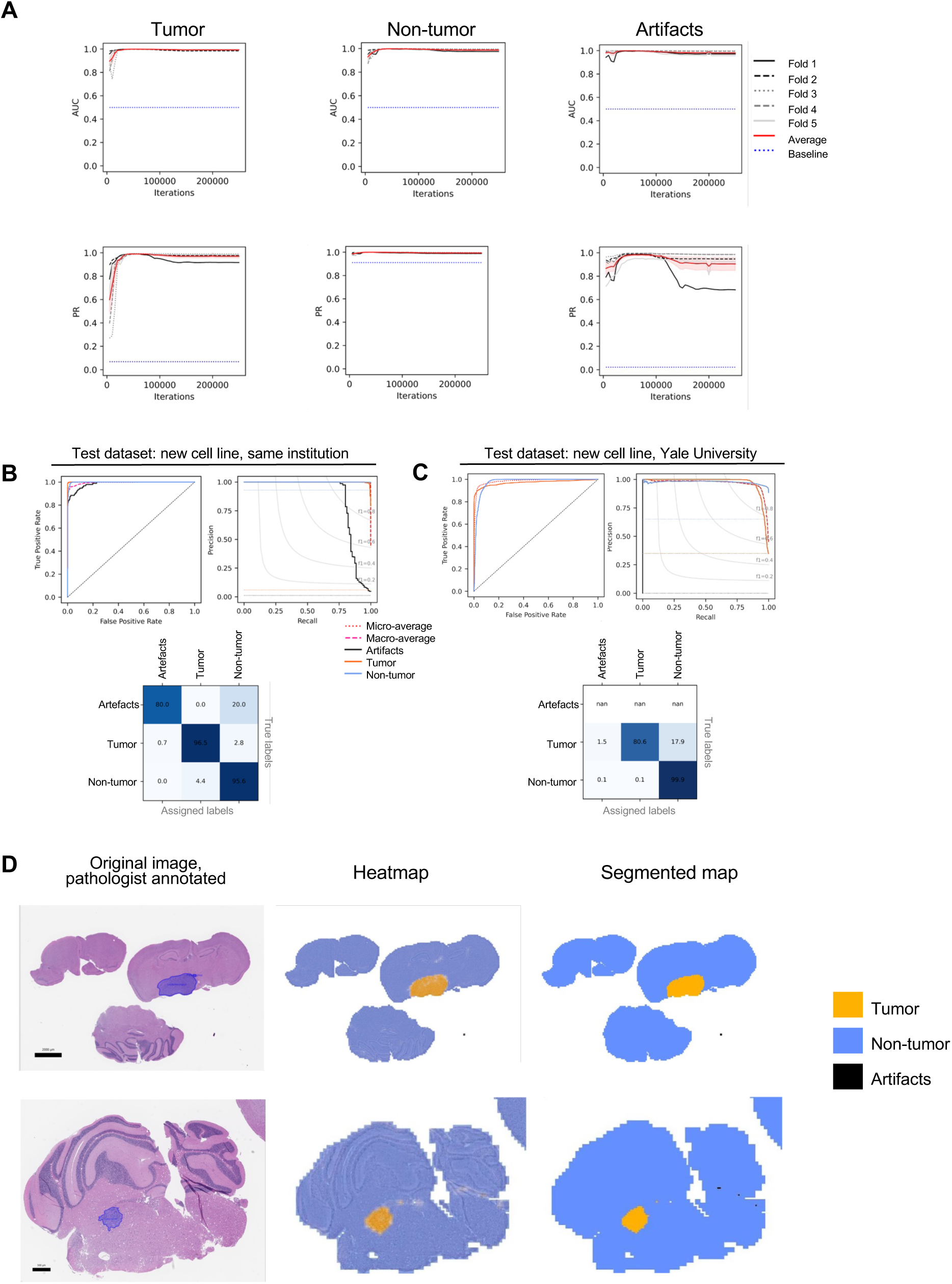
Identification of mouse brain metastatic tumors using deep-learning. (A) Evolution of the per-tile validation’s Area Under the Curve (AUC) and Precision Recall (PR) of the 5-fold training for each class. (B) Performance on the external datasets after training using 100% of the tiles. Top left: using the Receiver Operating Curve (AUC) curve and, top right: using the Precision Recall (PR) curve (in purple are shown the performances on the tumor region, in blue of the non-tumor tissue, and in plain black are shown the performance on artifacts annotations (for the ROC, the baseline performance of a random classifier is shown by the black dotted line; for the PR, the baselines performance appears in dotted lines of the same colors as the associated classes). Bottom left: confusion matrix. (C) Examples of whole slide images with manual annotation of tumor from pathologist (left), heatmap with tiles colored according to the majority class and intensity of the color proportional to the probability generated by the trained network (middle), and final segmented map with class assigned on which the tumor content can be extracted (right)

Nonetheless, a ROC of 0.971 is reached for detection of tumor (**Figure 2C**, confidence intervals CIIs=0.964-0.978) and 0.973 for non-tumor regions (CIs=0.968-0.978), and a PR of 0.969 and 0.977 for tumor and non-tumor (CIs=0.964-0.975 and CIs=0.970-0.983 respectively) (**Fig. 2C**). The confusion matrix shows an overall specificity of 0.938, an accuracy of 0.9543, and a precision of 0.6369 (only tumor and non-tumor annotations were available for this cohort) (**Fig. 2C**). **Figure 2D** shows representative images of segmentations done by our pathologist co-investigator and the neural network classifier on test cohort images.

### Training and validation of an AI-based liver metastasis identification tool

Similarly, H&E-stained liver sections from mice harboring xenograft melanoma metastases were annotated by a board-certified pathologist for 5-fold cross-validation training of a liver metastasis identifier. These training cohorts included 142 slides from 3 xenograft experiments, plus 16 sections from negative control mice (**Table S1**). The trained neural network distinguished tumor and normal tissue compartments within the liver with robust accuracy: on the 5-fold cross-validation, the liver metastasis classifier identified metastatic tumors with AUC of 0.978 (std=31e-3), normal non-tumor regions with AUC of 0.969 (std=56e-3), other tissues (e.g., blood, gastrointestinal tissue, muscle) with AUC=0.986 (std=35e-3) and artifacts with AUC of 0.979 (std=38e-3, **Figure 3A)**. For those same classes, the PR were respectively 0.979 (std=16e-3, baseline performance=0.043), 0.993 (std=9e-3, baseline=0.038), 0.850 (std=125e-3, baseline=0.019) and 0.872 (std=270e-3, baseline=0.020). The liver metastasis identifier algorithm was then tested on two external cohorts, including slides from xenograft and allograft models (**Figure 3B and C**). As summarized in **Table S6**, the model leads to AUC=0.998 (CIs=0.997-0.999) and PR=0.985 (CIs=0.988-0.990, baseline of 0.087) in detecting tumor, and AUC=0.981 (CIs=0.979-0.982) and PR=0.987 (CIs=0.985-0.988, baseline of 0.818) in detecting non-tumor from xenograft models, with specificity, accuracy and precision above 0.96 (**Table S7**). Though the training was not done with any slides from allograft models, the performance was still very high albeit slightly lower, with AUC=0.986 (0.969-0.999) and PR=0.978 (CIs=0.960-0.992, baseline of 0.068) for tumors regions, and AUC=0.941 (CIs=0.928-0.955) and PR=0.937 (CIS=0.920-0.953, baseline of 0.701) for non-tumor regions (**Table S6**), with specificity, accuracy and precision above 0.91 (**Table S7**). Due to the low number of annotations assigned to the “other tissues” class in the training and the high diversity of such tissue, it is not surprising that the network performance is lower for that category which is often mis-classified as non-tumor (confusion matrices in **Figure 3B-C**). However, given the rare occurrence of such tissues and the relatively small area they occupy when compared to the tumor and non-tumor regions, their impact on the final estimation of the tumor burden is unlikely to affect final estimates in most cases. As we collect more diverse samples in the future, we will be able to enlarge our database of “other tissues” and further improve these results. **Figure 3D** shows image examples from the test set, showing original regions from two xenografts and one allograft sample, and the corresponding heatmaps and segmented heatmaps obtained from the neural network classification pipeline.

**Figure 3.**
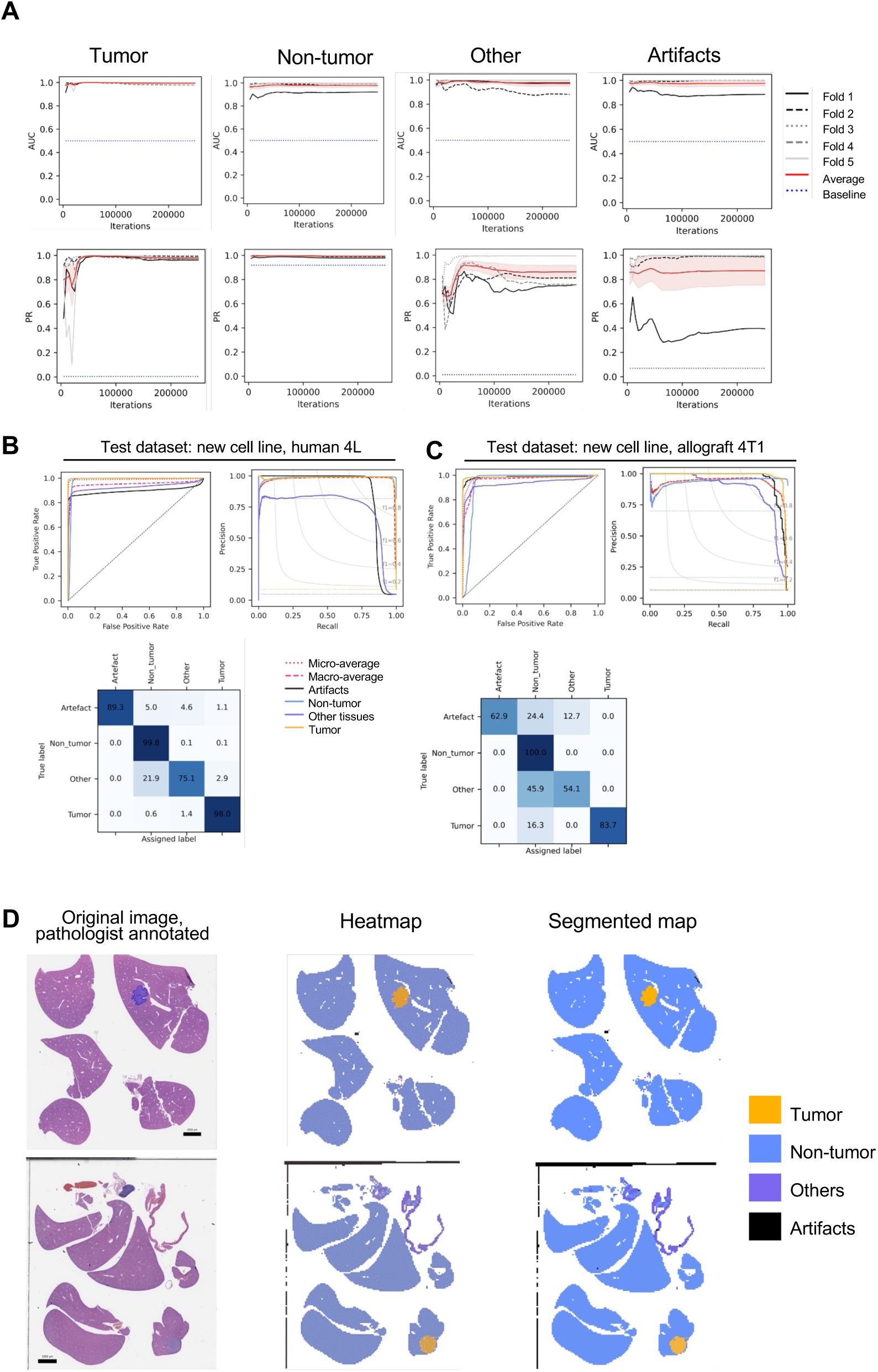
Identification of mouse liver metastatic tumors using deep-learning. (A) Evolution of the per-tile validation’s Area Under the Curve (AUC) and Precision Recall (PR) of the 5-fold training for each class. (B) Performance on the xenograft external dataset and. (C) Allograft external dataset after training using 100% of the tiles. Top left: using the Receiver Operating Curve (AUC) curve, and top right: using the Precision Recall (PR) curve (in purple are shown the performances on the tumor region, in blue of the non-tumor tissue, in purple of the other types of tissues, and in plain black are shown the performance on artifacts annotations (for the ROC the baseline performance of a random classifier is shown by the black dotted line; for the PR, the baselines performance appear in dotted lines of the same colors as the associated classes). Bottom left: confusion matrix. (D) Example of whole slide images with manual annotation of tumor from pathologist (left), heatmap with tiles colored according to the majority class and intensity of the color proportional to the probability generated by the trained network (middle), and final segmented map with class assigned on which the tumor content can be extracted (right).

### MetFinder: a web-based interface for tumor burden quantification

To make this tool easily and freely accessible to the preclinical cancer research community, we developed MetFinder, a web interface where scientists can upload their xenograft / allograft whole slide images and retrieve information such as tumor area, non-tumor area, tumor content (as a ratio of the tumor / non-tumor area), whole slide average tumor probability estimated by AI and tumor size distribution estimates (**Figure 4**).

**Figure 4.**
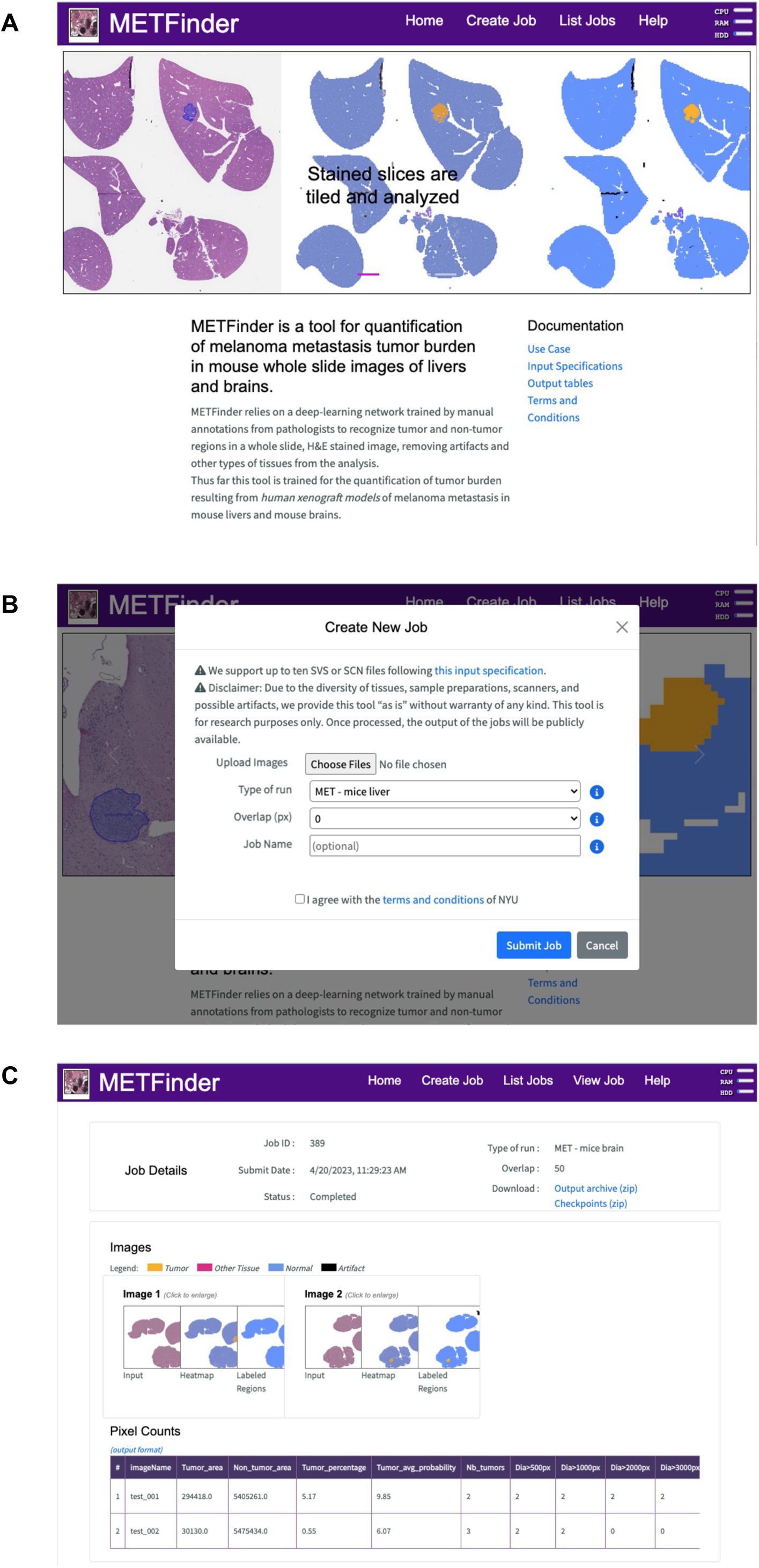
MetFinder interface. (A) The home landing page of the MetFinder site. (B) The dialog box where users can upload their images and specify which algorithm (liver or brain) they wish to use, as well as the degree of overlap between tiles. (C) The results page, where users can find the output of their job, described under Materials and Methods “MetFinder Web Interface”.

### MetFinder renders comparable results to manual annotation by an experienced experimentalist

To test the efficiency and accuracy of MetFinder in the hands of the end-user, tumor burden in livers as measured with MetFinder was compared to results obtained by manual annotation by a graduate student with histopathology training using OMERO Plus (Glencoe Software). We observed a high level of concordance between human annotation and MetFinder results (R^2^ = 0.9064, p < 0.0001; **Figure 5A**), when the tumor burden is rendered as a percentage of the total section area. Importantly, this strong correlation exists despite the fact that the tumor-bearing livers come from mice injected subcutaneously with a melanoma cell line that was not included in the training set, 501-MEL ^43^. An example of a hand-annotated liver section (left) is visible alongside MetFinder output (right) in **Figure 5B**. As reflected in the Pearson correlation, experimental results are consistent between methods. Critically, painstaking manual annotations of 29 sections took an estimated 6 hours of hands-on time by the student, who is proficient with OMERO and experienced in melanoma pathology. Using MetFinder, almost the same results are obtained with hands-on time limited to uploading the images. **Figure 5C, D** shows another liver dataset annotated by hand compared to MetFinder’s results. With R^2^ = 0.9828 and p < 0.0001, MetFinder’s output is again highly correlated with time-consuming annotations done by hand by an experienced researcher.

**Figure 5.**
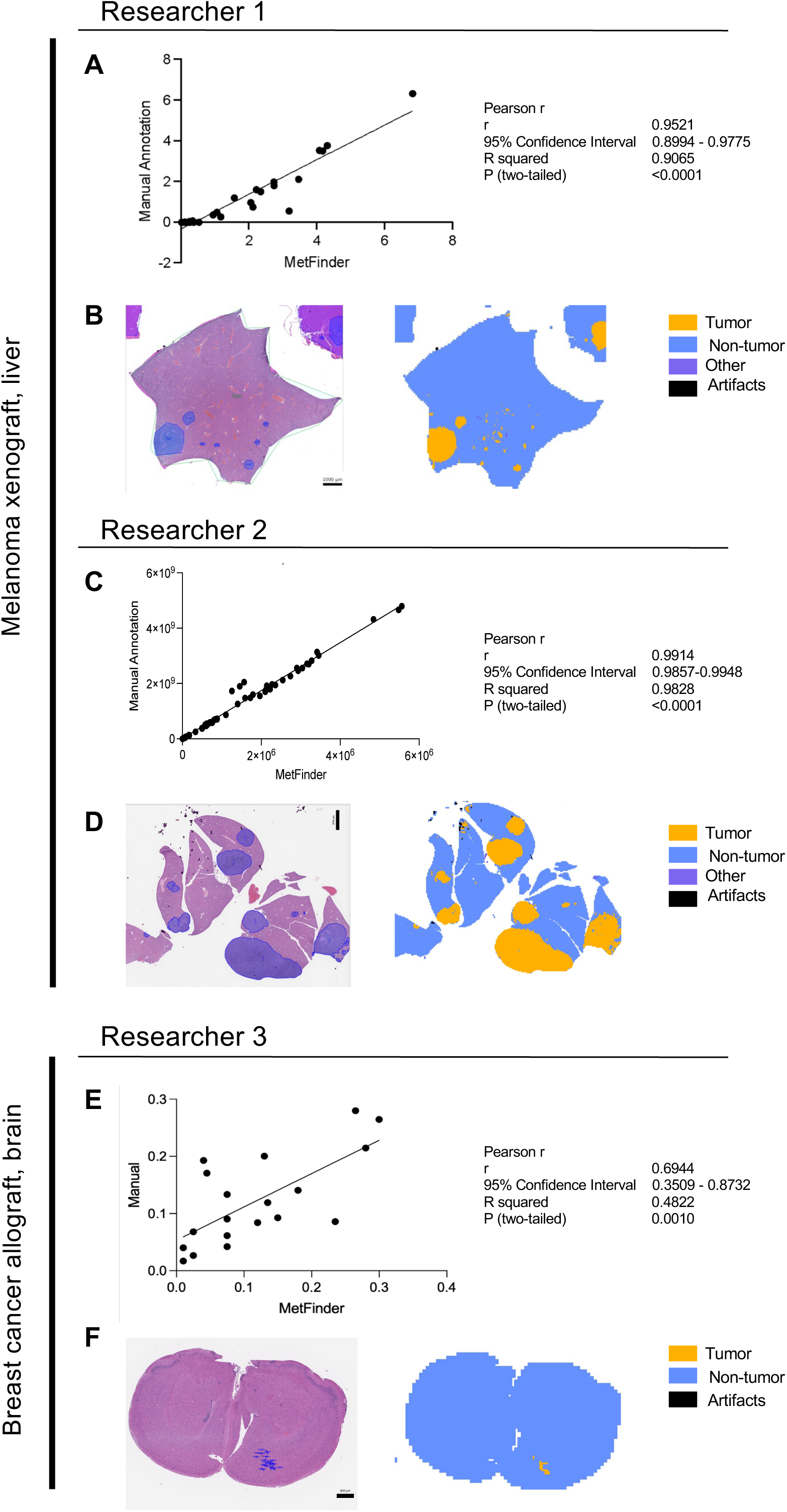
Comparison of AI-based tumor content estimation with manual annotation of tumor sections. (A) Correlation between manual annotation of tumor vs non-tumor area (represented as a percentage) by an experienced PhD student, representing a total of about 6 hours hands-on time, and annotation using MetFinder on 69 images from livers of 29 mice injected subcutaneously with the 501-MEL melanoma cell line. (B) Example of one part of one liver used in (A), hematoxylin and eosin stained input (left) and segmented output from MetFinder showing classification of each tile (right). (C) Correlation between manual annotation of tumor vs non-tumor area (represented in pixels) by a post-doctoral fellow, representing a total of about 8 hours hands-on time, and annotation using MetFinder on 69 images from livers of 22 mice injected subcutaneously with the MeWo melanoma cell line. (D) Example of one part of one liver used in (C), hematoxylin and eosin stained input (left) and segmented output from MetFinder showing classification of each tile (right). (E) Correlation between manual annotation of tumor vs non-tumor area (represented as a percentage) by an experienced PhD student, representing a total of about 6 hours of hands-on time, and annotation using MetFinder on 46 slides from brains of 19 mice injected intracardially with the allograft breast cancer line 4T1. (F) Example of one part of one brain used in (A), hematoxylin and eosin stained input (left) and segmented output from MetFinder showing classification of each tile (right).

In a further test of MetFinder’s capability, we compared its results with that of another manually annotated dataset (**Figure 5E, 5F**). The test was again applied to cells that MetFinder had not been trained to recognize. In this case, however, the tumors arose from a *breast cancer* allograft (4T1, ^44^ injected in a different mouse strain (BALB/c mice), a further biological divergence from the training cohort. Nonetheless the correlation with manual annotation was positive (R2 = 0.4822, p = 0.0010). This suggests that MetFinder is capable of delineating and analyzing tumors that arise from models other than melanoma.

### AI-based tools show comparable performance over other methods that quantify brain and liver metastatic burden

Next, we compared the ability of our AI-based Liver and Brain Metastasis identifier method to estimate metastatic burden compared to the most common currently used method, bioluminescence (BLI). 12273BM human melanoma cells transduced with a lentiviral construct encoding luciferase were injected intracardially into mice. The experiment was designed to test the hypothesis that a candidate gene (“Gene”) is essential to melanoma brain metastasis in an *in vivo* setting. Therefore, mice were divided into three groups of 12, with one group receiving cells expressing a control shRNA and two expressing a Gene-targeting shRNA. At the experimental endpoint, mice were injected intraperitoneally with luciferin and their metastatic burden was measured as previously described using the In Vivo Imaging System (IVIS, Xenogen Corp., Alameda, CA) ^42^ before euthanasia and organ harvesting. Once brain sections were H&E-stained and scanned, MetFinder was applied to also quantitate metastatic burden. The BLI data at termination are pictured in **Figure 6A**. To quantify tumor burden in the brain with BLI, a “Region of Interest” is drawn as an oval around the head of each mouse and the signal therein rendered as Total Flux in Living Image software (**Figure 6C**). When applied to brain H&E-stained sections (**Figure 6B**), MetFinder analysis of percent tumor burden reaches the same statistical conclusion by one-way ANOVA regarding the difference between groups in this experiment. That is, the knockdown of the candidate “Gene” renders decreased metastatic burden in the brain (**Figure 6D**). To quantify extracranial tumor burden by BLI, a rectangular ROI is drawn around the mouse body. Importantly, this signal may include sites of metastasis besides the liver, but the liver is nonetheless the main site of tumor burden and therefore we compared the extracranial BLI data (**Figure 6F**) to the MetFinder analysis of each mouse liver (**Figure 6E,G**). While the BLI data show stronger (lower p-value) differences between groups, the data reflect a similar trend of decreased metastasis in the Gene knockdown groups (**Figures 6F,G**). Importantly, these data illustrate that MetFinder may be used in conjunction with or in place of current gold-standard methods of quantification of tumor burden in a hypothesis-driven *in vivo* study.

**Figure 6.**
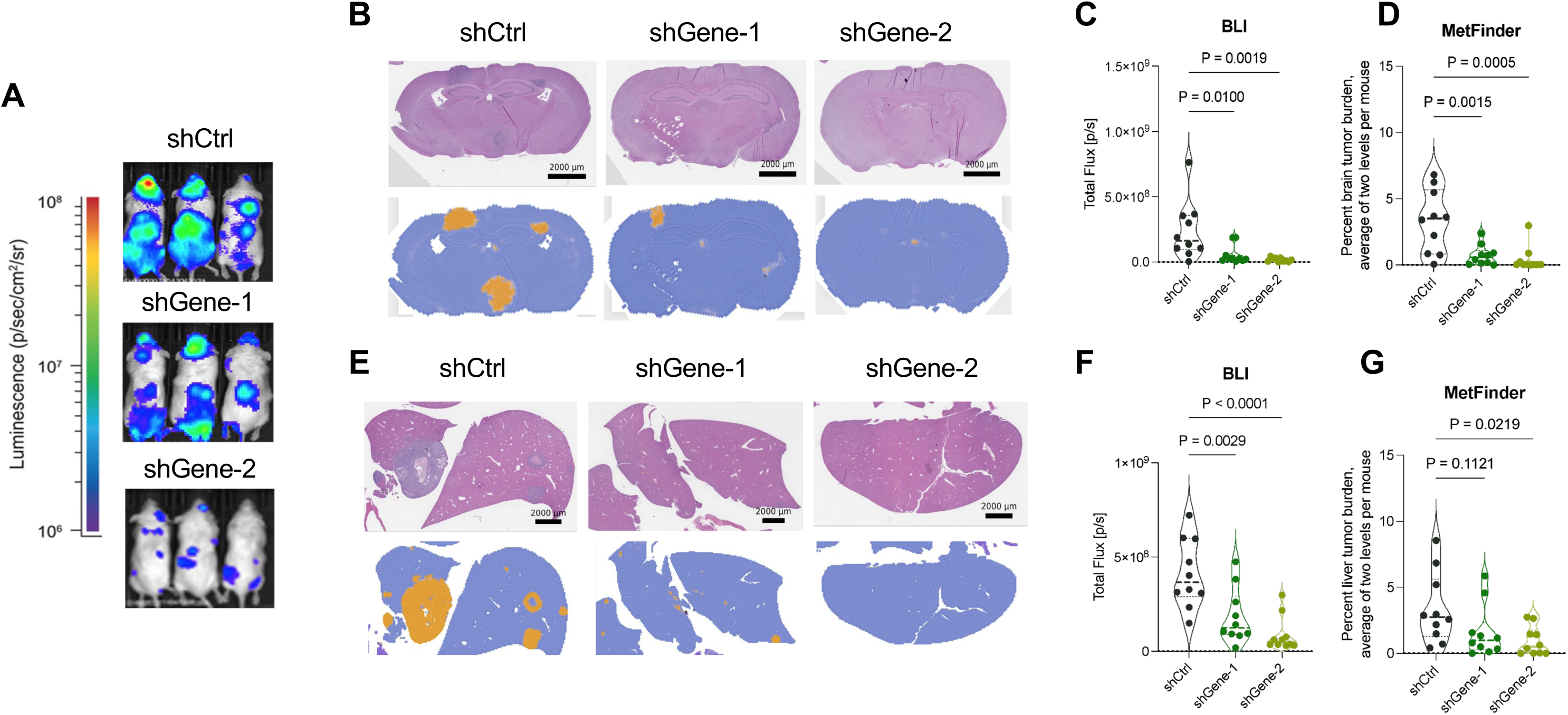
Comparison of AI-based tumor content estimation with bioluminescence imaging. (A) Representative bioluminescence images from 3 mice out of a total of n = 12 per group. Ctrl = non-targeting control shRNA. (B) Representative images from brains of 1 mouse per group; hematoxylin and eosin-stained input image (top) and final segmented map with class assigned on which the tumor content can be extracted (bottom). (C) Quantification of bioluminescence from a “region of interest” drawn over the brain of each mouse, avoiding the ears, eyes, and nose. Statistical comparison of groups by one-way ANOVA. (D) Total area of pixels assigned to tumor as a percentage of the total of tumor and non-tumor area for each brain slide were averaged for 2 slides (2 levels) per mouse. Statistical comparison of groups by one-way ANOVA. (E) Representative images from livers of 1 mouse per group; hematoxylin and eosin-stained input image (top) and final segmented map with class assigned on which the tumor content can be extracted (bottom). (F) Quantification of bioluminescence from a “region of interest” drawn over the trunk of each mouse, extending from the neck to the base of the tail. Statistical comparison of groups by one-way ANOVA. (G) Total area of pixels assigned to tumor as a percentage of the total of tumor and non-tumor area for each liver slide were averaged for 2 slides (2 levels) per mouse. Statistical comparison of groups by one-way ANOVA.

## DISCUSSION

Assessing metastasis burden in experimental *in vivo* models of cancer is crucial for understanding cancer progression, identifying potential therapeutic targets, and evaluating treatment efficacy. Several methods are used to assess metastasis burden in murine models. From lower to higher level of sensitivity and spatial resolution, Bioluminescence imaging (BLI), Positron emission tomography (PET), Microcomputed tomography (microCT), and Magnetic resonance imaging (MRI) allow monitoring metastatic dissemination non-invasively in live animals over time ^45^. These methods, however, are labor intensive and require access to expensive instrumentation. Post-mortem analyses, such as ex vivo BLI and ex vivo fluorescence imaging under a dissecting scope, provide additional information at termination of the experiment, but can be confounded by heterogeneity in reporter expression among tumor cells. Flow cytometry analysis on single cell suspensions of organs (using tumor surface markers), and quantitative RT-PCR of tumor-specific genes are also commonly used to estimate tumor burden on a given tissue ^9^, but fail to preserve information on morphological features and anatomical location. Ultimately, histological analyses (e.g., H&E, tumor-specific immunohistochemistry/IF stainings) are considered the gold standard for tumor load assessment, but require careful and comprehensive examination of multiple levels within a given tissue by an expert or trained experimentalist. This process is tedious, time consuming and subject to intra- and inter-operator variability.

We have developed an AI-based tool and corresponding online software (metfinder.org) to detect and quantify metastasis burden in sections of liver and brain tissues of mice carrying xenograft, allograft or syngeneic tumor implants. For that, we use a machine learning approach based on the DeepPATH pipeline, which has previously shown reliable results in classifying different types of human tissues ^29,46^, and is based on Google’s inception v3 architecture ^47^.

AI tools have shown promising results in assisting histopathological analyses, offering the potential to improve their accuracy, speed, and efficiency ^36,39^. These tools can help pathologists in various tasks, such as identifying and quantifying different tissue components, diagnosing diseases, and predicting disease outcomes ^25,27,29^. One of the significant benefits of using AI tools in histopathological analyses is the ability to process large amounts of data quickly and accurately. This capability can help reduce the workload and inter-observer variability, which is a common problem in histopathological analyses. By using consistent and objective criteria to analyze tissue samples, these tools can produce reliable and reproducible results.

Furthermore, MetFinder can provide valuable biological insights by yielding multiple layers of information about metastatic burden. These include both the relative area of metastasis and non-tumor organ parenchyma, as well as the number of metastases reaching various size thresholds. For instance, an experimental intervention which inhibits outgrowth of existing metastases but does not affect metastatic seeding may lead to many small tumors scattered throughout the organ. An intervention which affects seeding, however, may lead to fewer metastases of a size on par with the control. These observations will aid researchers in going beyond conclusions like “increased” or “decreased” metastasis to a more nuanced, mechanistic understanding of their experimental results.

Importantly, we have not tested the performance of MetFinder on genetically engineered mouse models, which differ morphologically from xenografts and allografts ^48^. Nor have we tested the ability of our tool to detect patient-derived xenografts, which are maintained in immunocompromised mice ^49^. Nonetheless, MetFinder displayed a remarkable ability to recognize melanoma metastases on tissue sections of mice injected with human and murine cell lines not initially included in the initial training (**Figures 2,3**), supporting the broad applicability of this tool. We used melanoma models as proof-of-principle for the ability to train a highly reliable AI tool, but observed that MetFinder also displays an impressive performance in detecting non-melanoma tumors (**Figure 4E,F**).

MetFinder requires uploading scans of histological sections to a server, which comes with potential limitations, such as the requirement for a stable and reliable internet connection. The quality and speed of the internet connection can significantly impact the user experience, because uploading of large images may be slow. In addition, MetFinder’s accuracy depends on the quality of the digital images uploaded to the server. Images that are blurry or have low resolution can lead to incorrect assessments, highlighting the importance of high-quality digital imaging equipment. Finally, differences in tissue processing (e.g., length and protocols of fixation and paraffin embedding), block sectioning, thickness and H&E staining, can introduce changes in tissue architecture and cell morphology that can impact MetFinder’s performance.

We recommend adhering as much as possible to the tissue processing and handling protocols reported in this article and the online tool to maximize MetFinder’s potential.

Overall, the use of AI tools in histological and pathological analyses shows great potential for improving accuracy, speed, and efficiency in in vivo experimentation for cancer research. With continued development and refinement of these tools, they have the potential to transform the way we analyze *in vivo* models. However, it is essential to note that AI tools are not meant to replace human expertise and judgment in histopathological analyses. These approaches should be viewed as complementary tools that can assist and support experts in their work.

## MATERIALS AND METHODS

### Experimental cohorts

Details regarding the datasets used in this study are summarized in **Tables S1-S3**. For the brain project, tissues from 3 different cell lines from 5 different intracardiac injection experiments were used, in addition to a cohort of negative control brains from mice that had not been injected: 4 for initial training and validation, (286 slides and 463,385 tiles total), and 2 as external test cohorts (71 slides generated from an experiment conducted at NYU leading to 8,536 annotated tiles and 18 slides leading to 4,521 annotated tiles for the cohort from Yale University). For the liver project, tissues from 6 different cell lines were used: 4 for initial training and validation (173 slides and 704,490 tiles total), and 2 as external test cohorts (58 slides with 208,910 annotated tiles and 14 slides with 3061 annotated tiles). The xenografted cell lines are derived from human melanoma metastases, the allograft from a genetically engineered mouse model (see **Table S3**). The cell lines were transduced with a lentivirus encoding a GFP-luciferase reporter and, in some cases, a genetic perturbation (CRISPR or shRNA). All mice were 8-12 weeks old at the time of intracardiac injection. At the experimental endpoint, mice were euthanized using humane protocols approved by our Institutional Animal Care and Use Committee (IACUC), and their organs were harvested.

### Tissue processing, sectioning and staining

Dissected tissues were briefly rinsed in cold PBS to remove excess blood and then immediately immersed in 10% neutral buffered formalin (Fisher Scientific cat# 4499), at a volume equivalent to 10-20 times the tissue volume. Tissues were fixed at room temperature for 72 hours with gentle shaking, rinsed 3 times in PBS and transferred to 70% ethanol. At this point brains were sectioned into thirds by coronal dissection with a razor blade. Livers are separated into lobes using a razor blade. These measures are taken in order to maximize the amount of organ surface area exposed in a single section. Samples were further dehydrated through graded ethanol and xylene, and then infiltrated with molten paraffin (Paraplast X-tra, Leica Surgipath cat# 39603002) in a Leica Peloris automated tissue processor. Paraffin processing times are provided in the **Tables S8 and S9**.

After processing, the tissues were embedded in paraffin blocks on a Leica Arcadia embedding system. The 3 parts of each brain were embedded such that the 1) olfactory bulb, 2) anterior half of the middle ⅓, and 3) posterior end of the last ⅓ are sectioned first. Livers are embedded such that all lobes are arranged in parallel. Five um thick sections were cut on a Leica RM2255 microtome with high profile disposable blades (Leica, cat#14035838926) and placed on superfrost slides (Fisher Scientific, cat# 22-042-924). Liver sections were taken from two distinct levels, separated by 100um. Brain sections were taken from three levels, each approximately one third through the depth of tissue, or ∼450um. Sections were deparaffinized and stained with hematoxylin (Leica cat# 3801575) and eosin (Leica, cat# 3801619) on an automated Leica ST5020 Multistainer. H&E staining times are provided in **Table S10**.

After the last xylene wash, slides were mounted with permount (Fisher, cat # SP15-500) and coverslipped (Leica Surgipath Premier Cover glass, 24×50, #1, cat #3800120ACS) on an automated Leica CV5030 coverslipper. The H&E-stained sections were scanned at a 40X magnification (pixel size 0.25 mm) on a Leica AT2 whole slide scanner and the image files uploaded to the NYUGSoM’s OMERO Plus image data management system (Glencoe Software).

### Data collection

The datasets were manually annotated at slide level by one of three board-certified pathologists (E.B., K.G. and G.J.) using either Aperio ImageScope (Leica Biosystems) or OMERO Plus (Glencoe Software). Annotation labels were “tumor”, “non-tumor”, “other tissues” or “artifacts”. For the brain, hardly any “other tissues” can be found and if so, they were grouped with the “artifacts” class. “Other tissues’’ include all non-relevant biological features that may have been retained from dissection, such as blood, gastrointestinal tissue, or muscle. Artifacts class, which can include pen marker, bubbles, or unknown objects, were supplemented with artifacts tiles from our database constituted with annotated tiles from various previous internal projects (32,730 tiles from 37 slides were added to the brain dataset, and 4,784 tiles from 32 slides were added to the liver dataset).

### Overall deep-learning approach

The overall workflow is described in **Supplementary Figure 2A** and detailed in the sections below.

### Training, validation and test process

The overall strategy is summarized in **Supplementary Figure 2A** and detailed below. The proposed tool is based on the pipeline developed previously ^23^ and based on the inception architecture ^41^. After annotations were done by the pathologists, the images were tiled at a magnification equivalent to 20x with a box size of 299×299 pixels (examples of tiles in **Supplementary Figure 2B**). Tiles were preprocessed using tile filtering (to remove tiles with no or little information, meaning tiles covered by more than 50% of background defined by values above an average gray level of 220 in 8 bits), color normalization using the Reinhard method ^50^. Also, a given tile was assigned a given manual label if its surface was covered by more than 50% by the mask drawn by the specialist. The training sets were split to achieve a 5-fold cross validation where the slides are split into 5 groups, and during each fold, 1 group is picked as the validation set while the other 4 are used for training. This cross-validation approach was used for optimization and to assess potential over-fitting, selection biases or potential variability inherent to such stochastic processes. Once done, the network fully re-trained with the whole train set was assessed with the external cohorts.

### Heatmap segmentation and tumor content estimate

To estimate the tumor content, we first generated heatmaps where each pixel is assigned a color associated with the label of the dominant class, and where the color is modulated by the probability associated with this class. If tiles were tiled with an overlap, the average probability is assigned to the corresponding pixels. To estimate the final tumor content, the heatmap needs to be thresholded, assigning a final label to each pixel corresponding to the dominant class (tumor, non-tumor, artifacts or other tissues). The surface of each class is estimated from the thresholded map. The tumor content is finally estimated as the ratio between the number of pixels labeled as “tumor” by the sum of pixels assigned either the label “tumor” or “non-tumor”.

### Performance evaluation

Several additional external tests were performed on slides from mice injected with different cell lines than the ones used during the training.

The first set of tests aim to assess the quality of the class assignment by the AI. A comparison of the manual tumor annotations and those resulting from the trained network was performed on a per-tile basis using the ROC (Receiving Operating Curve), the PR curve (Precision Recall) and statistics associated with the confusion matrix as implemented by the python scikit-learn library in multi-class configuration. To address how the potential uncertainties of annotations at the edge of a region (which may be imprecise or contain tissues associated with several labels), we studied those statistics first by assigning each tile the label of the mask if the mask covers at least 50% of the tile, and second by assigning each tile the label of the mask if the mask covers 100% of the tile.

Tests were then conducted to assess the effectiveness of the tumor content estimated by the AI, and the estimate generated by experienced scientists using manual annotation of the same images. Using OMERO Plus (Glencoe Software), users can draw a Region of Interest (ROI) on histological images. The area encompassed by a closed loop of any shape can be exported in a .csv file from OMERO. In the datasets used for comparison to manual annotation, the experimenters drew ROI’s for every tumor they found on each section, and also drew an ROI representing the area of the total organ parenchyma. The tumor burden can then be calculated as a percentage of the total area, just as in MetFinder’s output. Pearson correlations between MetFinder data and manual OMERO annotations were calculated using Prism 9.5.0, with a two-tailed t test.

Finally, we studied how tumor content compares with the most commonly used method of estimating tumor burden in mice, BLI. Imaging was carried out as previously described ^42^. Luminescence (Total Flux) within an oval ROI drawn around the head of each mouse was used to estimate brain tumor burden. Luminescence captured within a rectangular ROI drawn around the rest of the body was used to estimate extracranial metastasis. Using MetFinder, tumor content was estimated in two ways: 1) either using the percent of surface area occupied by tumor tissue (as extracted from the segmented heatmap), or 2) by using the tumor probability average on the whole slide using the heatmaps probability values before segmentation.

Pearson correlations between MetFinder data (% tumor area) and BLI (Total Flux) for each body region were calculated using Prism 9.5.0, with a two-tailed t test.

### MetFinder web interface

For screenshots of the interface at work please refer to **Supplementary Figure 1**. The outputs contain the heatmaps before and after thresholding and various measurements:

- The sum of pixels assigned the label “tumor”
- The sum of pixels assigned the label “non-tumor”
- The tumor burden, calculated as the ratio between tumor and sum of tumor and non-tumor regions
- The average tumor probability, calculated as the average of a tile being assigned the class “tumor” generated directly by the output of the network
- The estimated number of tumor sites, defined as the number of individual “objects” (sets of adjacent pixels) and as defined in the opencv library
- The estimated number of tumor sites distributions for various size thresholds
- The area in pixels of each tumor site (as estimated in opencv through the Green formula)
- The equivalent mean diameter of each tumor site (estimated by the average axis of the ellipse that best fits each tumor site, as defined by fitEllipse function from opencv)

The resolution of the final estimate will depend on the downscaling factor (set to 10 for the tests done in this manuscript and used to deal with memory limitations), and also the overlap of the tiles. With no overlap, the resolution is restricted by the 299316 cl:261299 pixel window, corresponding to a field of view of about 150316 cl:325150um.

## ACKNOWLEDGMENTS

We thank Dr. Iman Osman, Dr. Glen Merlino and Dr. Eva Perez-Guijarro for generously sharing patient-derived and murine melanoma cell lines. All embedding, sectioning, staining, and scanning were carried out by the Experimental Pathology Core at NYU Grossman School of Medicine. Experimental Pathology [RRID:SCR_017928] is part of the Division of Advanced Research Technologies and has received funds by the following grants: S10 OD021747 and Cancer Center Support Grant P30CA016087 at the Laura and Isaac Perlmutter Cancer Center. We thank Emil Rozbicki from Glencoe Software for help with image file interoperability using OMERO. The computational analysis for this work was supported by the NYU Langone High Performance Computing (HPC) Core’s resources and personnel. EH is supported by DoD Team Science Award-(W81XWH-20-MRP-TSA), P50CA225450 (NYULH Melanoma SPORE (PIs: Osman, Weber), R01CA274100, R01CA243446, R01CA277425 and U54CA2630001 (NIH/NCI; PIs: Hernando, Osman). N.E. is supported by CTSI (5TL1TR001447); G.B. was supported by T32CA193111; A.R. is supported by T32GM136542. The Research Software Engineering Core was supported by SCR_023022.

## AUTHOR CONTRIBUTIONS

A.K., N.C., A.T., and E.H. conceived and designed the study.

N.C. and A.T. applied neural networks and examined the performance of the algorithm.

E.B., K.G. and G.J. are board-certified pathologists who annotated sections.

A.K., A.A., S.S, N.E., A.Rubanov, M.N., R.M., G.B., G.L., and L.J. conducted in vivo experiments and provided tissue sections.

A.K., N.E., A.A.S.S, M.N., A.Rubanov, R.M. annotated sections.

V.M., M.A., C.L., processed tissues, sections, scanned slides and uploaded them to OMERO.

D.L. and A.Rubens generated and tested the MetFinder web interface.

A.K., N.C., A.T., and E.H. wrote the manuscript.

All authors read and approved the manuscript.

## DATA AND CODE AVAILABILITY

**MetFinder will be made freely available at** metfinder.org **for academic and non-commercial research use.** Requests for access to MetFinder may be directed to. Alternatively, to allow users to run the code on their own cluster, the code will be made publicly available at publication and is currently supplied as Supplementary Material with this manuscript. The checkpoints of the trained models are available upon request. Test images can be found at https://genome.med.nyu.edu/public/tsirigoslab/DeepLearning/MetFinder/slides.

---

**Supplementary Figure 1:**
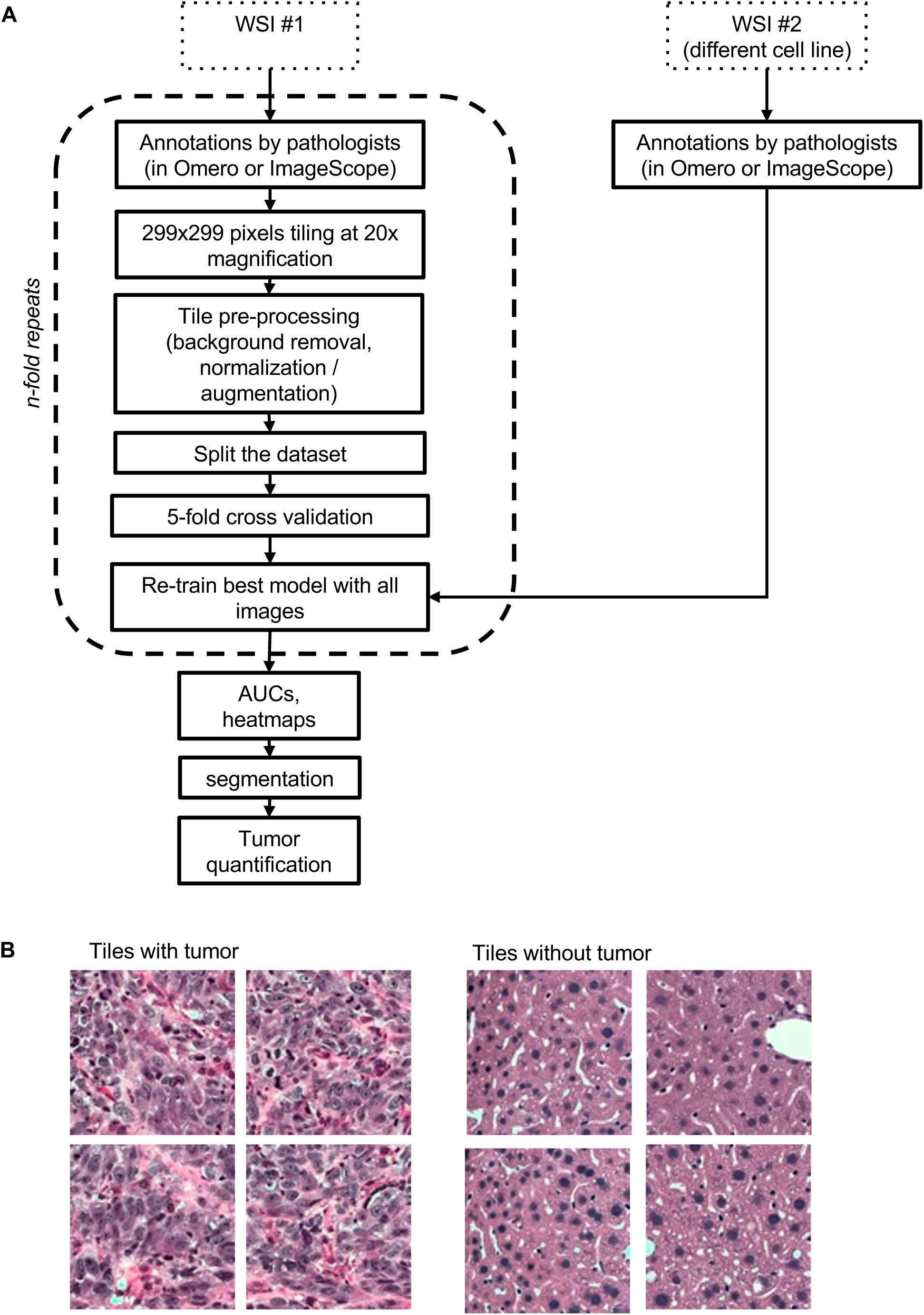
Development of a deep-learning tool to automatically characterize the size of tumors. (A) Development workflow. (B) Examples from mouse liver of 299419 cl:71299 pixels (about 150419 cl:93150 um) tiles at 20x magnification, which are used as a basis for learning to learn the labels

## SUPPLEMENTARY TABLES

**Table S1.** Details on the brain cohort used in this study.

| Test or Train | Source Institution | Dataset ID | Cell line | Xenograft or Allograft | Mouse background | Mouse sex | Number of slides | Number of mice |
| --- | --- | --- | --- | --- | --- | --- | --- | --- |
| Train | NYU | 20210111_AK | 12273BM | Xenograft | Nod/Scid/Il2gR null | Male | 70 | 35 |
| Train | NYU | 20210713_AK | 10230BM | Xenograft | Foxn1nu | Male | 91 | 36 |
| Train | NYU | 20220125_AK | 10230BM | Xenograft | Foxn1nu | Male | 70 | 35 |
| Train | NYU | 20220601_AK | 131/4-5B1 | Xenograft | Nod/Scid/Il2gR null | Male | 67 | 35 |
| Train | NYU | 20220927_AK | none | Xenograft | Nod/Scid/Il2gR null | Male | 14 | 8 |
| Test | NYU | 20211101_AK | 113/6-4L | Xenograft | Foxn1nu | Male | 71 | 35 |
| Test compare manual annotation Fig 4E-F | NYU | 20220928_NE | 4T1 | Allograft | BALB/c | Female | 46 | 19 |
| Test | Yale | 20230210_AK | YUGEN8 derivatives, YUKOLI and A375Br | Xenograft | Foxn1nu | Male | 20 | 20 |

**Table S2.** Details on the liver cohort used in this study.

| Test or Train | Source Institution | Dataset ID | Cell line | Xenograft or Allograft | Mouse background | Mouse sex | Number of slides | Number of mice |
| --- | --- | --- | --- | --- | --- | --- | --- | --- |
| Train | NYU | 20210111_AK | 12273BM | Xenograft | Nod/Scid/Il2gRnull | Male | 71 | 35 |
| Train | NYU | 20210511_GB | MeWo | Xenograft | Nod/Scid/Il2gRnull | Male | 9 | 9 |
| Train | NYU | 20210624_GL | SKMEL2 | Xenograft | Nod/Scid/Il2gRnull | Male | 62 | 31 |
| Train | NYU | 20220927_AK | none | Xenograft | Nod/Scid/Il2gRnull | Male | 16 | 8 |
| Test | NYU | 4L20211101_AK | 113/6-4L "4L" | Xenograft | Foxn1nu | Male | 58 | 35 |
| Test | NYU | 20211004_Allog | Mel114433 "M1" | Allograft | C57/Bl6 | Male and Female | 14 | 14 |
| Test compare manual annotation Fig 4A-B | NYU | C7_61_1079_AN R | 501-MEL | Xenograft | Nod/Scid/Il2gRnull | Female | 69 | 29 |
| Test compare manual annotation Fig 4C-D | NYU | 20210525_RM9036_CH391 | MeWo | Xenograft | Nod/Scid/Il2gRnull | Male | 69 | 22 |

**Table S3.** Details on the cell lines used in this study.

| Cell Line ID | Source |
| --- | --- |
| 12273BM | Dr. Iman Osman (NYU Langone) <sup>7</sup> |
| 10230BM | Dr. Iman Osman (NYU Langone) <sup>7</sup> |
| 131/4-5B1 | Dr. Robert Kerbel (University of Toronto) <sup>51</sup> |
| 113/6-4L | Dr. Robert Kerbel (University of Toronto) <sup>51</sup> |
| MeWo | ATCC |
| SKMEL2 | ATCC |
| Mel114433 "M1" | Gift from Dr. Glenn Merlino <sup>52</sup> |
| 501-MEL | Dr. Ruth Halaban (Yale University Medical School) |

**Table S4.** AUC and PR performances of each class for the brain metastasis classifier, assuming either 50 or 100% overlap between the test tiles and the annotated masks when assigning the labels.

|  |  | 50% mask/class overlap |  |  |  | 100% mask / class overlap |  |  |  |
| --- | --- | --- | --- | --- | --- | --- | --- | --- | --- |
|  |  | 2 tumor classes |  | tumor classes combined |  | 2 tumor classes |  | tumor classes combined |  |
|  |  | score | baseline | score | baseline | score | baseline | score | baseline |
| AUC | Leptomeningeal | 0.991 | 0.500 | 1.000 | 0.500 | 0.996 | 0.500 | 1.000 | 0.500 |
|  | Intrparenchymal | 0.971 | 0.500 |  |  | 0.984 | 0.500 |  |  |
|  | Non-tumor | 0.997 | 0.500 | 0.997 | 0.500 | 1.000 | 0.500 | 1.000 | 0.500 |
|  | Artefact | 0.986 | 0.500 | 0.986 | 0.500 | 1.000 | 0.500 | 1.000 | 0.500 |
| PR | Leptomeningeal | 0.757 | 0.043 | 1.000 | 0.059 | 0.866 | 0.035 | 1.000 | 0.042 |
|  | Intrparenchymal | 0.258 | 0.017 |  |  | 0.160 | 0.007 |  |  |
|  | Non-tumor | 1.000 | 0.930 | 1.000 | 0.930 | 1.000 | 0.938 | 1.000 | 0.938 |
|  | Artefact | 0.853 | 0.011 | 0.853 | 0.011 | 0.983 | 0.020 | 0.983 | 0.020 |

**Table S5.**
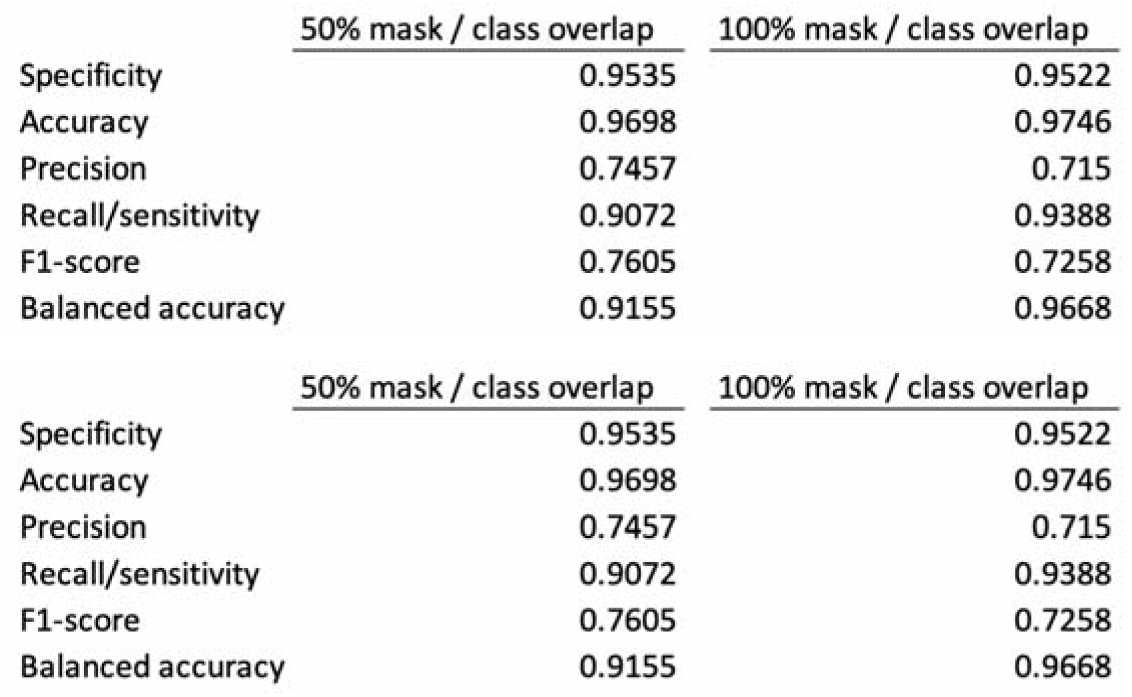
Performance of brain metastasis classifier on external cohort after slide segmentation.

**Table S6.**
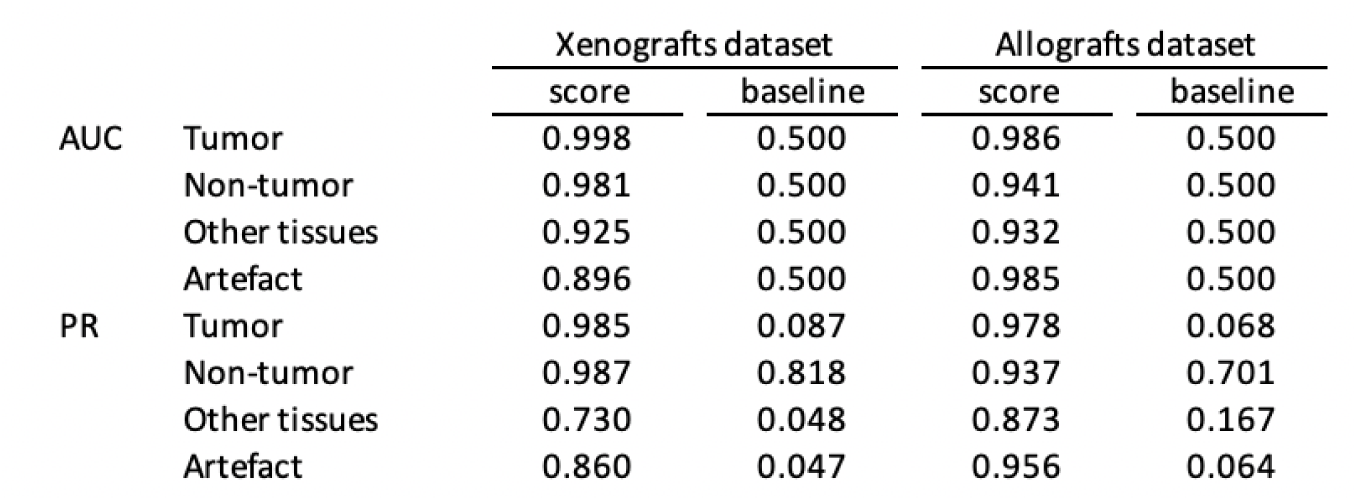
AUC and PR performances of each class for the liver metastasis classifier, using a 50 overlap between the test tiles and the annotated masks when assigning the labels.

**Table S7.** Performance of liver metastasis classifier external cohort after slide segmentation.

|  | <u>Xenografts dataset</u> | <u>Allografts dataset</u> |
| --- | --- | --- |
| Specificity | 0.9783 | 0.9113 |
| Accuracy | 0.9895 | 0.9443 |
| Precision | 0.9669 | 0.9471 |
| Recall/sensitivity | 0.9056 | 0.752 |
| F1-score | 0.9334 | 0.824 |
| Balanced accuracy | 0.9251 | 0.7086 |

**Table S8.** Paraffin processing times for liver tissues.

| Step | Reagent | Time (min) | Temp. | Pressure/Vacuum |
| --- | --- | --- | --- | --- |
| 1 | EtOH (70%) | 1 | Ambient | Ambient |
| 2 | EtOH (graded) | 1 | Ambient | Ambient |
| 3 | EtOH (graded) | 1 | Ambient | Ambient |
| 4 | EtOH (graded) | 1 | Ambient | Ambient |
| 5 | EtOH (graded) | 15 | Ambient | Ambient |
| 6 | EtOH (graded) | 15 | Ambient | Ambient |
| 7 | EtOH 100% | 30 | Ambient | Ambient |
| 8 | Xylene | 1 | Ambient | Ambient |
| 9 | Xylene | 10 | 45 | Ambient |
| 10 | Xylene | 35 | 45 | Ambient |
| 11 | Paraffin | 10 | 58 | Vacuum |
| 12 | Paraffin | 10 | 58 | Vacuum |
| 13 | Paraffin | 35 | 58 | Vacuum |

**Table S9.** Paraffin processing times for brain tissues.

| Step | Reagent | Time (min) | Temp. | Pressure/Vacuum |
| --- | --- | --- | --- | --- |
| 1 | EtOH (70%) | 1 | Ambient | Ambient |
| 2 | EtOH (graded) | 1 | Ambient | Ambient |
| 3 | EtOH (graded) | 1 | Ambient | Ambient |
| 4 | EtOH (graded) | 1 | Ambient | Ambient |
| 5 | EtOH (graded) | 20 | Ambient | Ambient |
| 6 | EtOH (graded) | 20 | Ambient | Ambient |
| 7 | EtOH 100% | 45 | Ambient | Ambient |
| 8 | Xylene | 1 | Ambient | Ambient |
| 9 | Xylene | 10 | 45 | Ambient |
| 10 | Xylene | 45 | 45 | Ambient |
| 11 | Paraffin | 10 | 58 | Vacuum |
| 12 | Paraffin | 10 | 58 | Vacuum |
| 13 | Paraffin | 40 | 58 | Vacuum |

**Table S10.** H&E staining times.

| Step | Reagent | Time |
| --- | --- | --- |
| 1 | 65°C baking | 25:00 min |
| 2 | Xylene (Fisher Scientific, cat# X3P1GAL) | 2:00 min |
| 3 | Xylene | 1:00 min |
| 4 | Xylene | 1:00 min |
| 5 | 100% EtOH (Decon Laboratories, cat# 2701) | 1:00 min |
| 6 | 100% EtOH | 1:00 min |
| 7 | 95% EtOH (Decon Laboratories, cat# 2801) | 1:00 min |
| 8 | 95% EtOH | 1:00 min |
| 9 | Water | 2:00 min |
| 10 | Hematoxylin (Leica cat# 3801575) | 2:00 min |
| 11 | Water | 0:10 min |
| 12 | Water | 2:00 min |
| 13 | Clarifier (Leica, cat# 3803598) | 0:15 min |
| 14 | Water | 2:00 min |
| 15 | Bluing (Leica, cat# 3802918) | 2:00 min |
| 16 | Water | 2:00 min |
| 17 | 95% EtOH | 1:00 min |
| 18 | Eosin (Leica, cat# 3801619) | 1:30 min |
| 19 | 95% EtOH | 1:00 min |
| 20 | 100% EtOH | 1:00 min |
| 21 | 100% EtOH | 1:00 min |
| 22 | Xylene | 1:00 min |
| 23 | Xylene | 1:00 min |
| 24 | Xylene | 1:00 min |
| 25 | Xylene | Exit |

