## Supplemental Material 1 for "MetFinder: a neural network-based tool for automated quantitation of metastatic burden in histological sections from animal models"

*MetFinder* is a tool for quantification of melanoma metastasis tumor burden in mouse whole slide images of livers and brains.

MetFinder relies on a deep-learning network trained by manual annotations from pathologists to recognize tumor and non-tumor regions in a whole slide, H&E stained image, removing artifacts and other types of tissues from the analysis. Thus far this tool is trained for the quantification of tumor burden resulting from human xenograft models of melanoma metastasis in mouse livers and mouse brains.


# Web access
MetFinder can be access at http://spawebwcdcdvm01.nyumc.org/. Slides can be downloaded via the webinterface and will run on our local computers.

# Local installation
If you want to run MetFinder on your own cluster machines, it will have to be run via command line.

First, install the DeepPATH pipeline via the [DeepPATH github page](https://github.com/ncoudray/DeepPATH/).

Then, you will need the checkpoints associated with the brain and those associated with the liver met (checlpoints are freely available to academic labs upon request).

The `sb_MetFinder.sh` script above is a template script written for slurm/bash clusters. See the  [DeepPATH github page](https://github.com/ncoudray/DeepPATH/) for more details about the different instructions. In the script above, you will need to change the variables declared in the first few lines:
- The `#SBATCH#` instructions are specific to our SLURM cluster. You need to adjust to whatever cluster you are using.
- you will need to instal the `conda3_520_env_deepPath.yml` environment, as described in [DeepPATH github page](https://github.com/ncoudray/DeepPATH/), and eventually adjust the `conda activate` line according to the path where your environement is created
- `PATH_TO_CODE` corresponds to the path where the DeepPATH is saved on your machine
- `PATH_TO_BRAIN_CHEKPOINTS` and `PATH_TO_LIVER_CHEKPOINTS` are the paths where the checkpoints downloaded previously have been saved
- `nbox` and  `noverlap` can be modified to `299` and `0` respectively if you want to run MetFinder without tile overlap (save time and space)

`sb_MetFinder.sh` expects 2 inputs: first the full path to the svs image to process, second the string 'LIVER' or 'BRAIN' depending on which algorithm you want to run. On cluster, several images within a folder can then be processed in parallel by initiating one for each of them. For example:
```shell
for f in `ls  path_to_svs_images/*svs`
do
	fff=`basename $f`
	echo $fff
	sbatch --job-name=job_name_${fff} --output=rq_liver_${fff}_%A.out  --error=rq_liver_${fff}_%A.err  sb_All.sh $f 'LIVER' 
done
```

Notes: 
- the architectures were trained on slides scanned at 20x on a Leica AT2 whole slide scanner, therefore leading to a pixelsize of above 0.5 um/pixel. 
- the sample code requests 1 GPU but the code should run on CPU alone too (though slightly slower).

The ouputs will be saved in the folder in which the command is run. Within subfolders carrying the original name of each image, you should find the same 3 jpg images as those provided on the website (original tiles analyzed, the heatmap with intensity of the colors proportional to the associated probability, and the final segmented map. In the heatmap and the segmented map, tumors will appear in orange and normal tissue in blue, with artefacts in black and other types of tissues in purple). You will also find 1 csv with the different measurements done:

-   `Total tumor area` and `non-tumor area`: total number of pixels assigned each label, at 20x (most scanners have a pixel size around 0.5 um / pixel at this magnification).
-   `Tumor_percentage`: Tumor_Area / (Tumor_Area + Non-Tumor_Area)
-   `Tumor_avg_probability`: The AI assigned each tile a probability of belonging to a certain class (tumor, non-tumor, artifacts, other). The probability of all tiles is averaged and given here.
-   `Nb_tumors`: Total number of tumors (estimated as individual sets of non-connected tumor regions)
-   `Nb_tumors_500px_Dia_or_more`, `Nb_tumors_1000px_Dia_or_more`, etc …: Total number of tumors with a diameter above 500px, 1000px, etc…
-   `List_of_tumor_areas`: detailed list of all measured tumor areas
-   `List_of_tumor_diameter`: detailed list of all measured tumor diameter (estimated by the average of the min and max axis)


# Template slides
Template slides can be downloaded from (https://drive.google.com/file/d/1629CycCv1JAss5EhnwDSbfzLaWkE612O/view?usp=share_link)
